# Food Webs: Insights from a General Ecosystem Model

**DOI:** 10.1101/588665

**Authors:** Cesar O. Flores, Susanne Kortsch, Derek Tittensor, Mike Harfoot, Drew Purves

## Abstract

Food webs have been intensively studied throughout modern ecology, using empirical evidence, statistics and modelling tools to search for consistent patterns, underlying commonalities, and variations between food webs and ecosystems. However, with few exceptions, the modelling approaches have not been based on the emergent properties of complex simulated ecosystems. For the same reason, there have also been few studies of ‘sampling the model’, in which different levels of sampling effort are imposed in a controlled simulated environment to explore the effects of varying sampling intensity in order to relate those back to empirical observations. Here, we introduce the Madingley Model, a general ecosystem model based on ecological and biological first principles of interactions between individuals, as a potential tool for analyzing food webs. In doing so, we present the first insights of the analyses of emergent Madingley food web networks. We describe the basic structure of these networks and introduce a process to aggregate and sample the food webs produced by this model so that it can be compared to empirical food web studies. We show that food webs created by this model reasonably reproduce the properties of empirical food web networks. Furthermore, we provide insights about the effects of species aggregation and sampling on the observed structure of empirically documented food webs.

## Introduction

The rate and spatial scale at which humans are impacting the environment through activities such as agriculture, fisheries, clearing of forests, habitat fragmentation, pollution, and climate change, is unprecedented in history (Sala et al. 2000, Duraiappah et al. 2005, Levin and Lubchenco 2008), and is causing substantial alterations in ecosystem structure and functioning (Barnosky et al. 2012). Impacted ecosystems can have reduced resilience, be less complex, have less recovery potential, and be more prone to potentially irreversible phase-shifts or trophic cascades (Estes et al. 2011, Kortsch et al. 2012), with concomitant consequences for the supply of goods and services on which humans depend. To mitigate impacts and inform policy decisions about diversity and conservation, it is crucial to improve our understanding of how ecosystems work. Empirical studies and experiments play an important role; however, given the necessarily limited spatial extent and practical limitations for manipulative experiments, and the inherently large number of variables that cannot be controlled, they must necessarily be complemented by additional approaches to help disentangle key mechanisms and processes. Mechanistic ecosystem models, which simulate key communities and ecological interactions and then let ecosystem structure and function evolve, are one such tool for gaining insights into the mechanisms underlying ecosystem dynamics and structure by simulating key biological processes. Moreover, these models are useful frameworks for making forecasts about anthropogenic impacts on ecosystems and can be used to investigate trade-offs between various management options (Christensen and Walters 2004, Fulton et al. 2011, Purves et al. 2013, Harfoot et al. 2014).

Modelling ecological phenomena of specific ecosystems and addressing management questions with mechanistic multispecies models is not new (Christensen and Walters 2004, Krinner et al. 2005, Fulton et al. 2011). However, few have attempted to model the emergent properties of ecosystems arising from the basic interactions of simulated individuals, and instead tend to aggregate and specify mechanisms at higher levels. The Madingley model is the first General Ecosystem model (GEM), attempting to model emergent ecosystem structure and function across all marine and terrestrial environments globally by identifying, coupling and simulating the key biological processes (e.g. feeding, metabolism, reproduction, dispersal and death) of interacting individuals. One advantage of doing this is that higher level properties, such as food webs, assemble dynamically and can then be compared to empirical data for both assessing the ability of the model to capture observed patterns, and for gleaning further understanding of the processes that result in empirical observations of food webs.

The structural properties of food webs are emergent properties of interacting species in an ecosystem. Although dynamic and constantly shifting in terms of flux rates of matter and material and composition, food web properties tend to be studied as temporal averages or in case of topological approaches as binary linkages. A strength with the Madingley model is that it can be used to capture both the relative fluxes among entities, i.e. the species, as well as the binary structure of these food webs. Empirical food webs are known to exhibit emergent patterns and regularities, for example, the degree distribution (i.e. a measure of connectivity), the number of trophic levels, and the average path length (Dunne et al. 2002, Williams et al. 2002). The Madingley model can also be used to construct simulated food webs and use these to calculate metrics for comparison with empirical food webs. Although the Madingley model is based on individual interactions for the purposes of computational feasibility, individuals that share the same traits and follow the same ecological trajectories are aggregated together into **cohorts** (see Figure S1). These cohorts are the basic unit of the Madingley model. The simulations need a large number of cohorts in order to let the complexities of real ecosystems evolve and to provide sufficient coverage of trait space. Therefore, the aggregation into large numbers of cohorts (~1,000) result in simulated food webs where the amount of cohorts is 1-2 order of magnitude larger than empirical food web studies, where the number of species is of the order of ~10-100 (Dunne et al. 2002). We reduced the size of the Madingley networks to ensure appropriate comparisons between simulated and empirical food webs by developing cohort aggregation and a link sampling method in this study.

If the Madingley model can reproduce realistic food web structures, this provides a basis for further validation of the dynamic outputs from the model. Moreover, simulated food webs can be used to provide unique insights into how structural properties such as connectance might change along spatial and temporal gradients on a global scale, and how food web structure could change if an ecosystem is undergoing alterations due to environmental impacts such as species invasions or range shifts. Finally, the effects of sampling food webs at different intensities can be mimicked within the model system to help understand consequences of sampling effort for estimated metrics. To date most studies on sampling effort have been performed by analyzing incomplete empirical food webs (Winemiller 1990, Goldwasser and Roughgarden 1997, Martinez et al. 1999), while few studies have analyzed sampling efforts in model systems (but see Frund 2016). The main objective of this study has been to provide the first set of simulated food webs produced with a large-scale mechanistic ecosystem model, the Madingley model. We use the generated output firstly to extract conclusions about the validity of the model when compared to empirical food-web networks, and secondly to evaluate the output in light of aggregation and sampling methods in food webs.

## Materials and Methods

### Food webs based on the Madingley model

Food webs are representations of species and their feeding relationships (Yodzis and Winemiller 1999). In this study, we created food webs based on output simulations from the Madingley model. In our simulated food webs the species analogues are aggregations of individuals (cohorts) sharing the same set of traits, where individuals are grouped by either trophic similarity or body mass and size. We use food web network terminology and refer to these aggregations as nodes or “trophospecies”, whilst the feeding relationships between them are referred to as links. The Madingley model is a process-based, mechanistic modelling framework (Purves et al. 2013, Harfoot et al. 2014), in which ecosystem properties emerge from interacting individuals. During each time step, biological processes and interactions of cohorts (i.e. groups of individuals that are characterized by the same set of traits) through time and space occur. The basic processes that are modelled for each cohort are feeding, predation, metabolism, reproduction, and dispersal. Growth is an emergent property of the balance between metabolic expenditures and assimilated food material. The cohorts belong to 10 different functional groups, each with one or more different functional trait, such as diet (herbivores, omnivores, and carnivores), metabolic pathway (endotherm or ectotherm), reproductive strategy (semelparous or iteroparous). Each functional group is represented in the model by a set of cohorts with the same categorical traits by with potentially very different body masses. The model is forced with empirical and reanalyzed data on climate (Harfoot et al. 2014). For terrestrial ecosystems, a model of autotrophs determines vegetation structure based on climatic conditions and then allocates growth to leaves and structural tissues (Smith et al. 2013).

## Cohort aggregation

### Aggregation methods

To compare simulated food webs to empirical ones, it was necessary to aggregate cohorts based on their properties. We aggregated cohorts into ***S*** nodes or “trophospecies” (see step 1.1 in Figure 1) using two different approaches from the literature (Martinez 1991, Gilljam et al. 2011).

**Figure 1:**
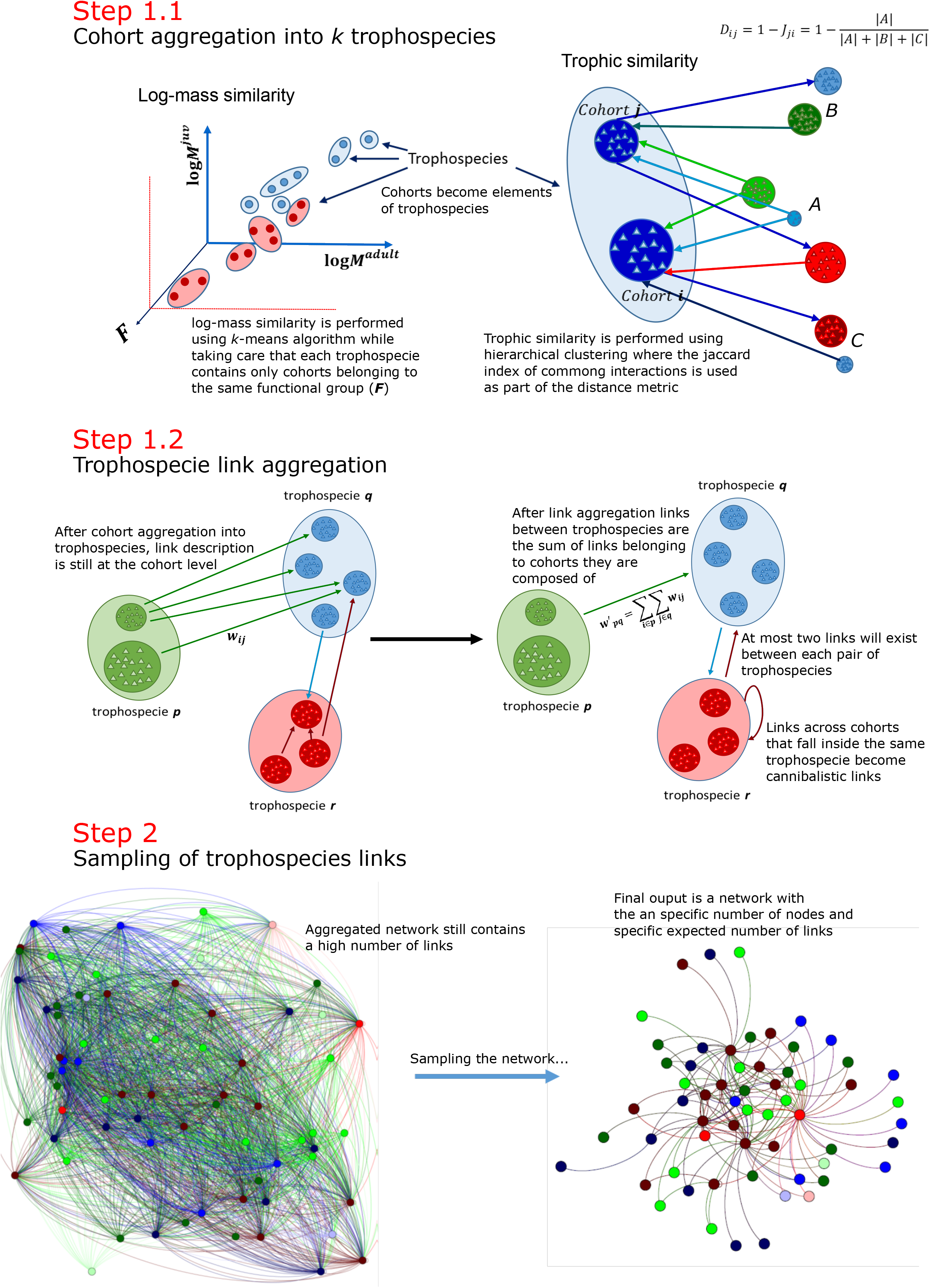
Flow of the Madingley food web aggregation and sampling. The full Madingley food web is first aggregated into *S* trophospecies using one of two different aggregation methods (step 1.1). During this step trophospecies are formed with cohorts belonging to the same functional groups, and properties of trophospecies are recalculated based in the cohorts that form them (Equations 2–4). As immediate step 1.2, links are also aggregated according to trophospecies they belong to (Equations 5), with links appearing if at least a link exists between cohorts they are composed of. Finally, as last step and in order to make the number of links comparable to those of empirical networks, links are sampled according to a Poisson Process described in Equations 6–8.

The first aggregation method is based on the **logarithmic mass similarity** between individuals proposed by (Gilljam et al. 2011). This approach clusters “species” or nodes in a food web according to their size or mass. To aggregate cohorts, we applied a ***K*-means** clustering algorithm (MacQueen 1967) based on Euclidean distance, to cluster along dimensions of logarithmic adult and juvenile mass of each cohort. We start the algorithm by randomly placing ***K=S*** centroids across the two dimensions following by an iterative process of ***(i)*** assigning cohorts to the centroid with the least squared Euclidian distance, and ***(ii)*** updating the centroid positions to the mean of the cohorts positions it contains. The iterative process is continued until convergence is achieved (i.e. assignments of cohorts to centroids do not change across iterations). Although final assignment depends on the initial random placing of centroids, results presented here were robust across different runs.

The second method is based on **trophic similarity** proposed by (Martinez 1991) and is commonly used. This method involves performing a **hierarchical clustering** based on trophic overlap between clusters (within functional groups). Trophic overlap can be described as the number of common interacting trophospecies between the two trophospecies over the total number of links they share, which is the Jaccard similarity index between trophospecies *i* and *j*:

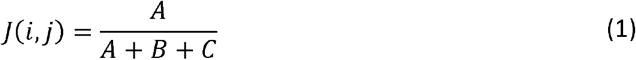

where *A* is the number of common predator and prey species that both *i* and *j* have, and *B* and *C* the number of predators and prey exclusive to *i* and *j*, respectively. Values range between 0 (no common predators or prey) and 1 (identical set of predators and prey). Initially each cohort is assigned to a different trophospecies. We group pairs of trophospecies according to the lowest Jaccard distance (**1** –*J*(*i,j*)). We repeat this process until we end up with ***S*** trophospecies. When two trophospecies contain more than one cohort, we use the average of the Jaccard distance between every pair of cohorts inside both nodes.

After cohorts are aggregated to their respective trophospecies, links between two trophospecies will prevail as long as at least one link exists between the original cohorts they are composed of. Because a trophospecies may be composed of cohorts with links between them, cannibalistic links will appear at this stage (see step 1.2 in Figure 1).

### Trophospecies property readjustment

Once trophospecies are aggregated from their initial cohorts, their properties are readjusted based on the cohort properties (see Figure S1) they are composed of. During this process we keep both the number of individuals and biomass constant before and after aggregation (i.e. both quantities should be independent of the aggregation method). Finally, the individual mass is readjusted as a weighted average. This process is summarized by the following equations:

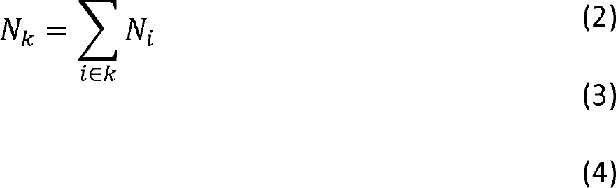

where *N_k_, B_k_*, and *M_k_* represent the number of individuals, biomass, and individual mass for trophospecie ***k***, respectively.

The same goes for biomass fluxes (link weights). New biomass fluxes between trophospecies need to consider all biomass fluxes between their constituent cohorts. By following this method, any interactions between cohorts inside the same nodes are represented by ‘cannibalistic links’ (see step 1.2 in Figure 1), where a link points from the prey to the predator and its weight *W_ij_* is the amount of biomass ingested by unit of time. Therefore, new biomass flux 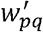 across two trophospecies *p* and *q* can be represented as:

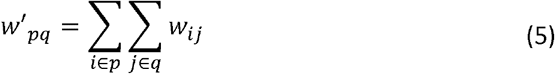

Where *W_ij_* represents the original biomass flux (mass by unit of time) in the original output of a Madingley simulation between a prey and a predator cohort.

### Link Sampling

Food webs generated by the Madingley model are highly connected in comparison to empirical networks. The main reason of the high connectance is that during a simulation we can observe all interactions, no matter its importance for the development of either prey or predator involved in them. By contrast, in nature these interactions it is difficult to observe all interactions for several reasons, including the rarity or some interactions and that they occur for small sized organisms or in unobservable locations. So, it can be hard to include or discard them. In order to remove “non-important” links, we sampled the network by assuming that encounters between prey and predators are governed by a Poisson process. Although these encounters cannot entirely be described using a homogeneous Poisson process (Beyer and Nielsen 1996) (i.e. individuals are not randomly distributed in space), our original assumption can still be used for the purposes of this article (i.e. to have a method of sampling the networks that has some resemblance of what happens in nature).

The Poisson process in our case will represent the times at which we will observe encounters between prey *p* and predator *q*, which can be described as a counting process *N*(*τ*) that represents the number of observed encounters up to a time *τ*. Then, the probability of observing *n* encounters in a time *τ* is given by:

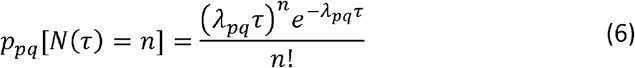

where *λ_pq_* represents the rate by unit of time at which encounters between predator and prey happens. In reality the last quantity depends on many factors (i.e. the sampled area, the number of observed individuals and by consequence the population size of each species). However, under our sampling approach we assume that it depends only on the biomass flux *w*′_*pq*_. The probability of keeping the encounter *p* → *q* depends then on the probability of observing at least one encounter up to a time *τ*:

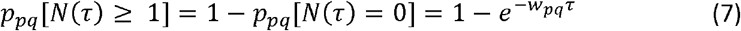

with *τ* (observation time) becoming the parameter that controls the strength of the sampling (sampling effort). The full set of probabilities between each pair of trophospecies at a specific observation time (*τ*) can be represented as an adjacency probability matrix **P**(*τ*) of size *S × S* where rows and columns represent prey and predators, respectively. We can then create a **sampled food web** with a binary adjacency matrix **A**, by applying the following element-by-element binary operation:

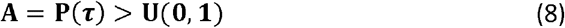

where **U**(**0,1**) is a random matrix with each element being a uniform random number between 0 and 1. One property of Equation 7 is that given infinite observation time (*τ*), the probability of observing at least one encounter becomes a certain event for any link *p* → *q* for which the biomass flux 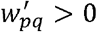.

### Network metrics

To characterise the structure of the simulated food webs, we focused on both **local** (e.g. no. of basal species) and **global** (e.g. modularity) food-web properties that are commonly measured in food webs (Dunne et al. 2002, Yen et al. 2016). All metrics were applied to the binary (as opposite to weighed) adjacency matrices of the sampled food webs (Equation 8). For **local** properties, we evaluated the **number of basal** (herbivores) *N_B_*, **number of intermediate** *N_l_* (with both prey and predators), and **number of top** *N_T_* (carnivores) trophospecies. Furthermore, **generality** *L* / (*N_l_* + *N_T_*) (average number of incoming links per predator), and **vulnerability** *L* /(*N_l_*+ *N_B_*) (average number of outgoing links per prey) were also included as part of our local metrics. Among the **global** properties we included the (i) **mean path length** *(i.e.* the mean path distance between every pair of connecting trophospecies, (Williams et al. 2002), the (ii) **clustering coefficient** (*i.e.* the probability that two neighbours of a trophospecies are neighbours themselves, (Watts and Strogatz 1998, Girvan and Newman 2002), and the (iii) **modularity** *(i.e.* which is a quantitative value between 0 and 1 describing how densely sub-groups of species interact with one another compared to species from other sub-groups (Newman and Girvan 2004). **Mean trophic level** (i.e. the shortest path from the base to each species) and **mean level of omnivory** (i.e. the level of omnivory of each species is the standard deviation of the trophic level of its resources) are global metrics that were also included in the analysis. The **connectance** *(i.e.* the fraction of observed links over all potential links *L* / (*S*S*)) was used to assess the dependence of food-web metrics on networks with different number of nodes.

### Empirical Data

We compared the simulated food webs with 52 empirical food webs (see empirical food webs listed in Table S1 and S2) using the aforementioned network metrics. In order to have a more direct comparison to empirical food webs, aggregation and sampling was performed using the Madingley output to match each of the empirical networks presented this study. For each study we aggregated the Madingley output to match the number of nodes *S* of the corresponding study. Then, we sampled the aggregation 1,0 times using Equation 8 where *τ* was chosen such that the expected number of observed links *E[L]* will match the one of the corresponding study. We then compared and evaluated the simulated network metrics against the empirical ones.

## Results

### Sampling effort

An important driver of food web patterns is the observation time *τ* (sampling effort). Figure 2 and S3 show the effect of sampling effort on an aggregated food web of *S = 60* trophospecies using *K*-means mass similarity and Jaccard trophic similarity aggregations, respectively. We can see qualitatively that the network pattern varies as we increase the sampling effort. Of special interest are the intermediate values of sampling effort, in which the well-known triangular pattern of food webs expressed as an adjacency matrix emerges when *K-means* mass similarity is used to aggregated cohorts, but not when Jaccard trophic similarity is used. In contrast, the latter is the most similar to the original pattern of a full Madingley food web (see Figure S2). Another well-known food web metric affected by the sampling effort is the degree distribution (Dunne et al. 2002). Figure 3 and S4 show that the expected degree distribution for an aggregated food web of *S* = 200 trophospecies using both methods, and a full Madingley food web, respectively. The shape of the degree distribution may be indicative of how well a food web has been sampled, with a uniform shape being oversampled and a power-law shape being under-sampled.

**Figure 2:**
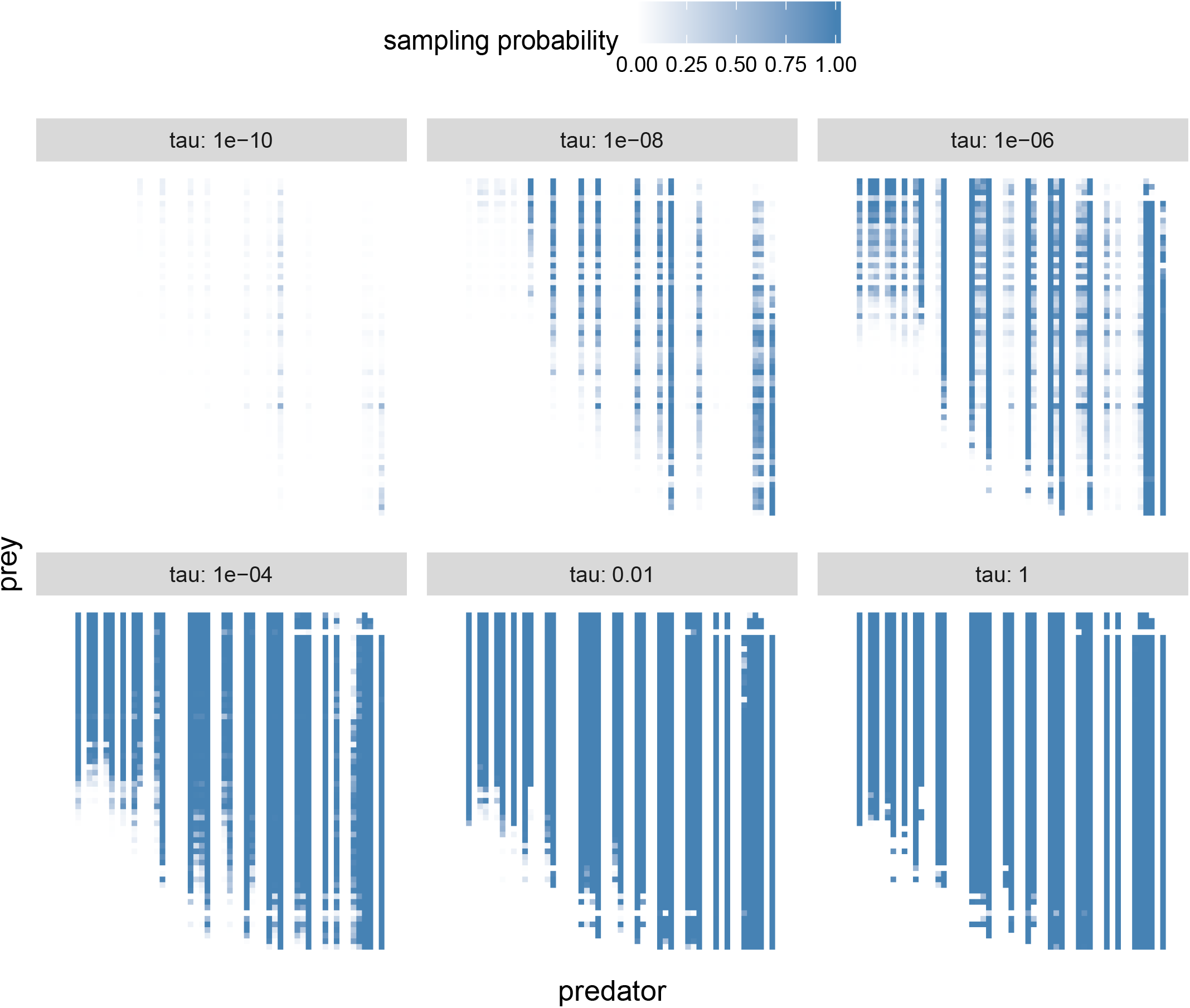
Effect of Sampling on an aggregated food web using *K-*means aggregation method. The figure shows the sampling probability of observing an interaction between preys and predators on an aggregated Mandingley food web, represented as an adjacency matrix for different observation times (sampling effort) *τ′s*. Cohorts were aggregated into *S* = 60 different trophospecies using the ***K*-means** aggregation method. Rows (prey) and columns (predator) are sorted according to increase individual mass.

**Figure 3:**
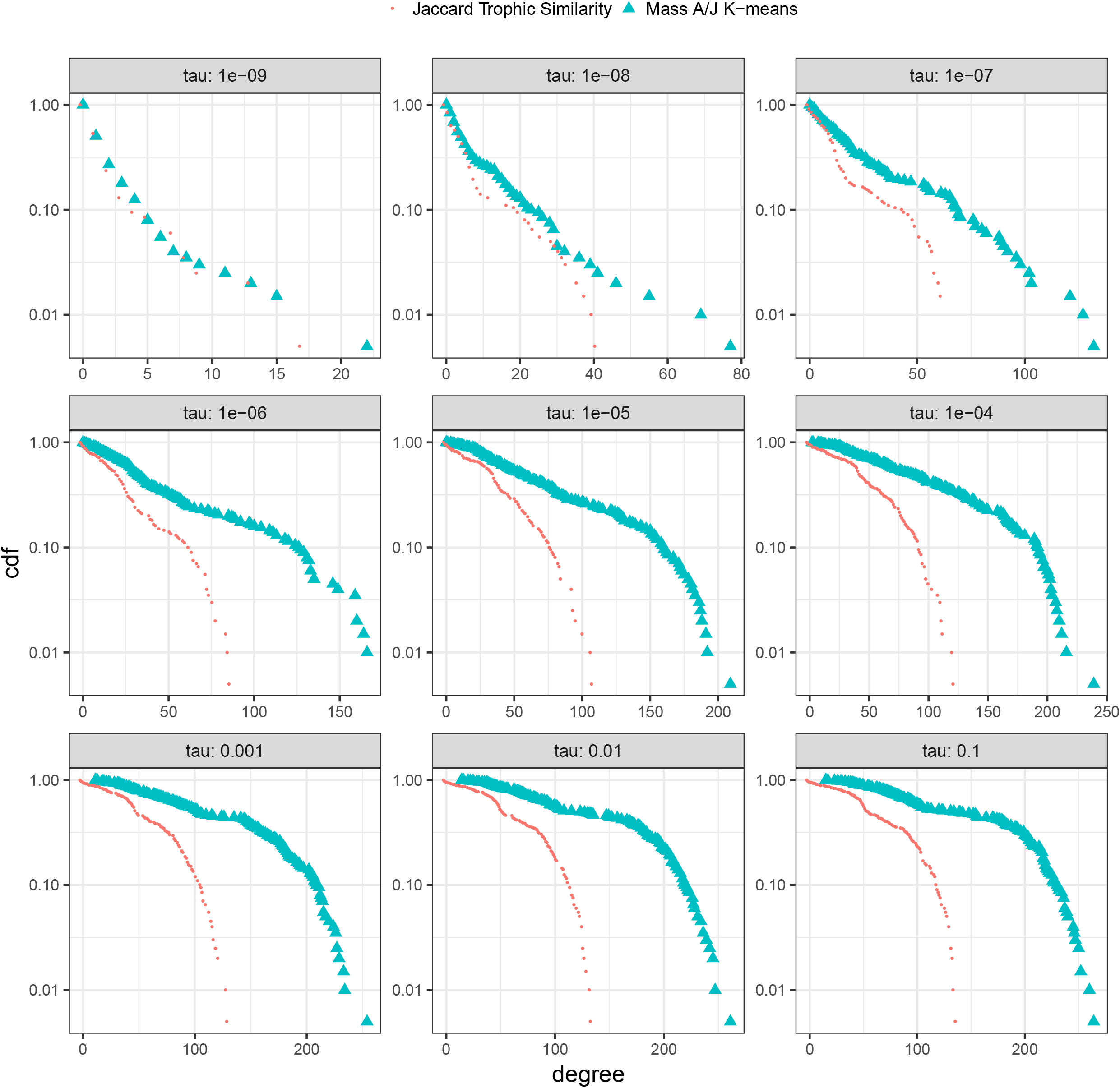
Effect of sampling on the degree distribution of Madingley Aggregated networks. The figure shows the expected degree distribution given observation time (sampling effort) *τ* for aggregation of the full Madingley food web into *S* = 100 trophospecies for the two different aggregation methods. Form of the degree distribution depends on both aggregation method and sampling effort. Furthermore, Jaccard aggregation method gives in general a smaller number of links (i.e. Sampling in Jaccard aggregated food webs should be stronger than *K*-means aggregated food webs in order to have the same expected connectance).

### Aggregation and sampling strategy effect on network metrics

As Figure 4 shows, there exists a dependency of network metrics on (i) aggregation methods. First, aggregating by trophic similarity produces higher values for both modularity and mean path length than aggregating by mass similarity, independent of the connectance value. Modularity result may be explained by the fact that these this metric directly involves the presence of adjacent links between trophospecies, and therefore becomes higher-valued in an aggregation method that benefits from the presence of trophospecies with cohorts that share the same links.

**Figure 4:**
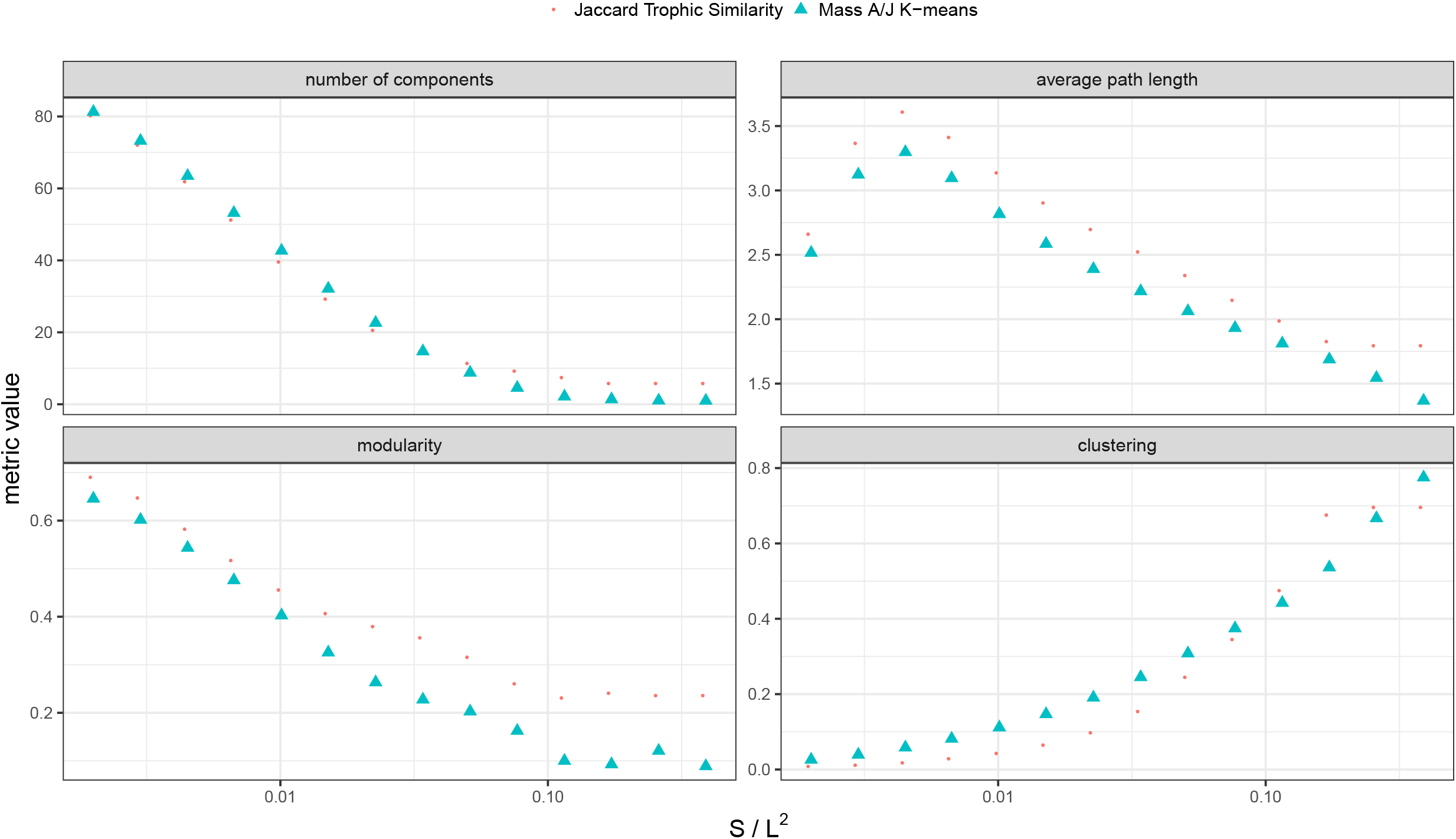
Effect of sampling and aggregated method different network metrics. The figure shows plots for different network metrics for the two aggregated methods. Each data point represents the mean of 1,000 aggregated food webs of *S* = 100 trophospecies. Sampling parameter τ was chosen to match the the corresponding connectance of each data point.

### Relationship between simulated and empirical food web studies

How dissimilar (mean deviations) or similar (values close to zero) the simulated Madingley food webs are to their corresponding empirical food webs depends on the metric considered (Figure 5, and S5). Generally, deviations between the outputs of the simulated and the empirical food webs fall into four classes. The first class are those metrics that tend to show generally small deviations between the Madingley simulated food-web and the empirical food web metrics such as the vulnerability, the number of top predators, mean level of omnivory and mean trophic level. The second class of metrics, for example the number of basal and intermediate species, are those metrics that display deviations as either negative or positive trends with increasing connectance between the simulated and empirical food web values. Another feature of this class is that the higher the connectance value, the more the simulated and empirical food web values converge. The third class consists of metrics that are scattered around the zero value, displaying both positive and negative deviations, such as modularity, without any clear bias or trend. Finally, the fourth class consisting of generality and clustering metrics, we find that the Madingley networks have, generally, higher values than empirical networks. Furthermore, for modularity and mean path length we can observe that empirical values can be in general larger or smaller depending on the aggregation method that was chosen for creating the simulated food webs.

**Figure 5:**
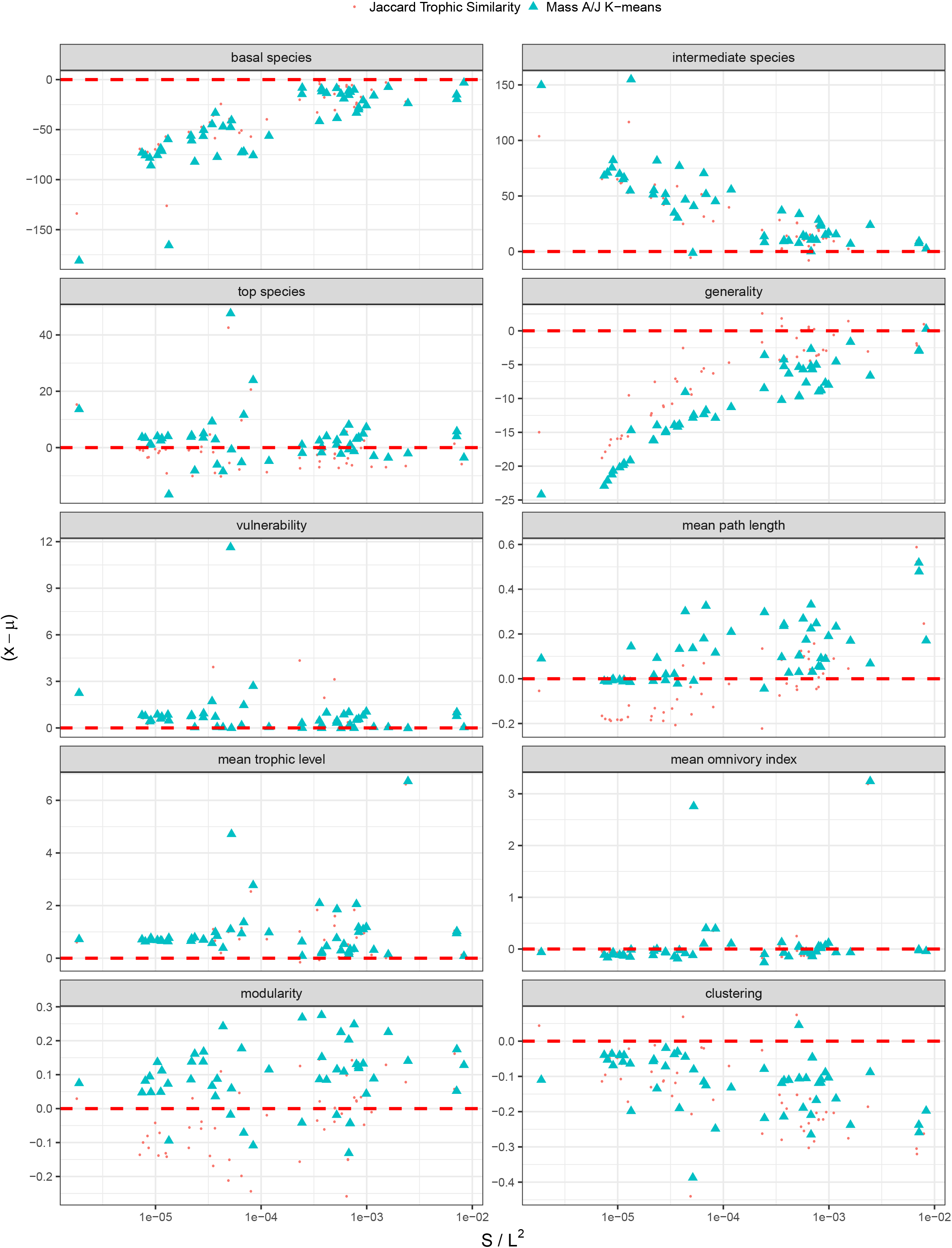
Comparison of Madingley sampled networks with empirical studies. For each empirical study 1.0 aggregated sampled food webs were created with the same number of species and expected number of links (i.e. same expected connectance) by choosing the appropriate parameter sampling effort ***τ***. Two data points (one for each aggregated method) exists for each empirical network, y-axis represent the absolute difference between empirical values ***x*** and the average of the corresponding Madingley sample food webs ***μ**.*

## Discussion

Here we present the first analyses of simulated food webs produced by a general mechanistic ecosystem model, the Madingley model. Our study explores how sampling effort impacts network structure and properties of a food web. The sampling effort in food webs is known to affect food web properties such as the number of links per species (Goldwasser and Roughgarden 1997, Riede et al. 2010). The shape of degree distribution in empirical studies does not display universal behavior and appears to depend on connectance and food web size (Dunne et al. 2002). Furthermore, our study using different levels of sampling (see Figure Figures 3 and S4) indicates that shape of the degree distribution on a food web may also depend on how well a food web has been sampled. Should that be the case, deeper study of the degree distributions produced by models such as Madingley could be an important tool on inferring missing links of empirical studies. Our study also shows that already at intermediate sampling effort the characteristic triangular shape of food web matrices emerges, which implies that at a given point adding extra sampling time/effort does not yield additional changes in food web structure. This is good news for empirical ecologists since building empirical food webs is time-consuming.

Finally, we also show how the way in which cohorts are aggregated into trophospecies can have a strong effect on the way the network structure appears (see Figures 5 and S5). Of special interest are modularity and clustering (two metrics where local neighbors structure is important), where aggregating by Jaccard trophic similarity will have in average a higher value than aggregating by K-means mass similarity. This is happening because the former aggregation method is using local network structure (i.e. neighbors) to perform the aggregation, while the latter does not use network structure information.

Our analyses also show that the Madingley model captures realistic features of empirical food webs such as trophic levels and level of omnivory, suggesting that the model can mimic emergent structural properties of ecosystems and that first-order ecological processes (e.g. vulnerability, mean trophic level, mean omnivory index on Figure 5) drive these patterns. However, for other metrics, the fit between simulated and empirical food web outputs depends on the chosen metric, the connectance value and the sampling effort. Examples of these metrics are basal and intermediate number of species as well as generality on Figure 5. In those metrics we observed that the difference between the Madingley food webs and empirical studies decreases as the connectance value increases.

Some metrics showed relatively little deviations between the modelled food webs and the empirical ones e.g. the vulnerability metric (i.e. predators per prey), suggesting that that this descriptor may equally well (or poorly) captured in both empirical and simulated food webs. The number of basal, intermediate and the generality metrics also showed an interesting pattern in which values of the empirical and simulated food webs converge as connectance increases. This convergence pattern of a better fit between the modelled and the empirical approaches for highly connected food web could reveal either sampling biases in the empirical food webs or aggregation problems in our modelled food web. For empirical food webs, scaling relationships between food web descriptors have been frequently reported (Riede et al. 2010, Digel et al. 2014), for example number of species scales inversely with connectance, implying that smaller (i.e. low number of species) food webs are more highly connected than larger ones (Riede et al. 2010). In addition, the fraction of basal decreases with number of species, whereas the fraction of intermediate species tends to increase (Martinez et al. 1999)

It is well-known that the number of basal, intermediate and top species in empirical food webs are strongly dependent on observational effort in the field (Martinez et al. 1999). As a result, these estimates are often biased towards better resolution at higher trophic levels and poorer resolution at lower trophic levels (Kortsch et al. 2015). In addition, diet information for low trophic level species, such as invertebrate, is often less resolved than for top predators (Kortsch et al. 2015). For the relatively low-connected food webs in our analyses, the simulated food webs produced food web networks with a higher fraction of basal species/nodes than the empirical ones. With respect to the deviation in the fraction of basal species between the simulated and empirical food webs for low-connected networks, the simulated food webs may represent a more realistic food web structure than their empirical counterparts, because empirical webs have historically underrepresented the number of basal species due to highly aggregated groups (e.g. phytoplankton) at lower trophic levels (Polis 1991, Kortsch et al. 2015). The proportion of basal, top and intermediate species are confounded. For example, the less basal species in a network, the more intermediate and top species and vice versa (Digel et al. 2014, Wood et al. 2015). Therefore, also with respect to the deviation in the fraction of intermediate species, the simulated food webs may give a more realistic output than the empirical, as the fraction of intermediate species most likely is overestimated and the fraction of basal species the underestimated in *large* empirical food webs. Most interestingly, the fraction of top predators is similar for both simulated and empirical food webs, suggesting that this feature may be more accurately represented in both empirical and modelled approaches. Indeed, in empirical networks, information on species at higher trophic levels is often more detailed and better resolved compared to information about species at lower trophic levels (Kortsch et al. 2015). The observed deviation between the simulated Madingley food webs and the empirical food webs predict well the anticipated bias of number of basal, number of intermediate and number of top species in empirical food webs, lending support to the chosen aggregation methods for the simulated food webs in this study.

Mean trophic level showed a general positive deviation, except for a few large outliers, indicating that the Madingley model tends to produce lower values than those from empirical studies. This could be due to the Madingley food webs containing more basal species at lower trophic level, yielding a lower mean trophic level for the whole network.

Not all metrics showed a relatively unbiased fit, for example clustering. This metric was studied for food webs by Dunne et al (Dunne et al. 2002), in which it was suggested that food webs, in general, lack this property, at least for small network sizes, but for larger food webs and other networks, clustering increases as a power-law of the number of nodes. This fact contrasts with the Madingley food webs, where clustering is on average consistently larger than empirical studies, indicating neighbors of nodes are more likely to be connected. This fact may suggest that the Madingley model can still be improved by incorporating additional traits in each cohort that further separate what a cohort can prey from (instead of the present mass-based relationship). For example, diurnal variation in activity levels may separate nocturnal and daytime predators and prey or having additional habitat specificity in terms of predator and prey affiliations. Adding these extra traits may decrease clustering of the networks to make them more similar to values observed in reality.

All of the insights gleaned from this work come from analyzing output from dynamically-assembled and emergent food webs produced by the Madingley model. We only considered static snapshots of the food webs; however, a deeper understanding of how food webs assemble and evolve may come from analyzing the dynamic data from the model. This model can allow us to perform simulations in order to test recent ideas about network stability (Allesina and Tang 2012, Serván et al. 2018). Ultimately, the model and empirical data can be used in an iterative synthesis; the empirical data to inform the model and validate its ecological properties, and the model to propose hypotheses and detect mechanisms that can then be tested with empirical data. We propose that, ultimately, the development of such models may provide a new and useful tool to assist with food web ecology.

