## Supplementary figures and tables for "Food Webs: Insights from a General Ecosystem Model"

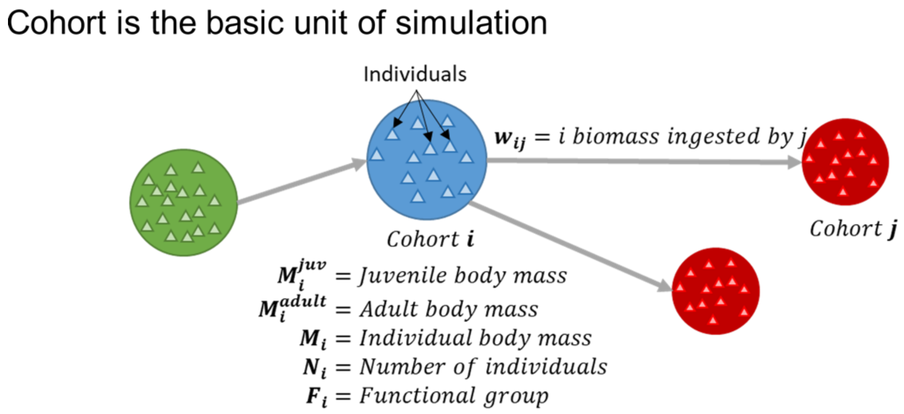


**Supplementary Figure 1: Cohort as the basic unit of simulation on the Madingley Model**. A Madingley simulated food web composed of cohorts and links that represent predator-prey relationships in terms of biomass transfer rates pointing from prey to predators. A cohort is defined as an entity with a certain number of individuals, each one with the same set of properties. For the analysis presented on this article, 24 snapshots at different times of a single run were used. Each snapshot represents a Madingley simulated food web composed of ***S = 1,000*** cohorts grouped into 9 different functional groups. Furthermore, each snapshot is composed of approximately ***L ≅ 160,000*** links, where the corresponding weight represents the biomass flux ($w_{ij}$) that goes from prey ***i*** to predator ***j***.

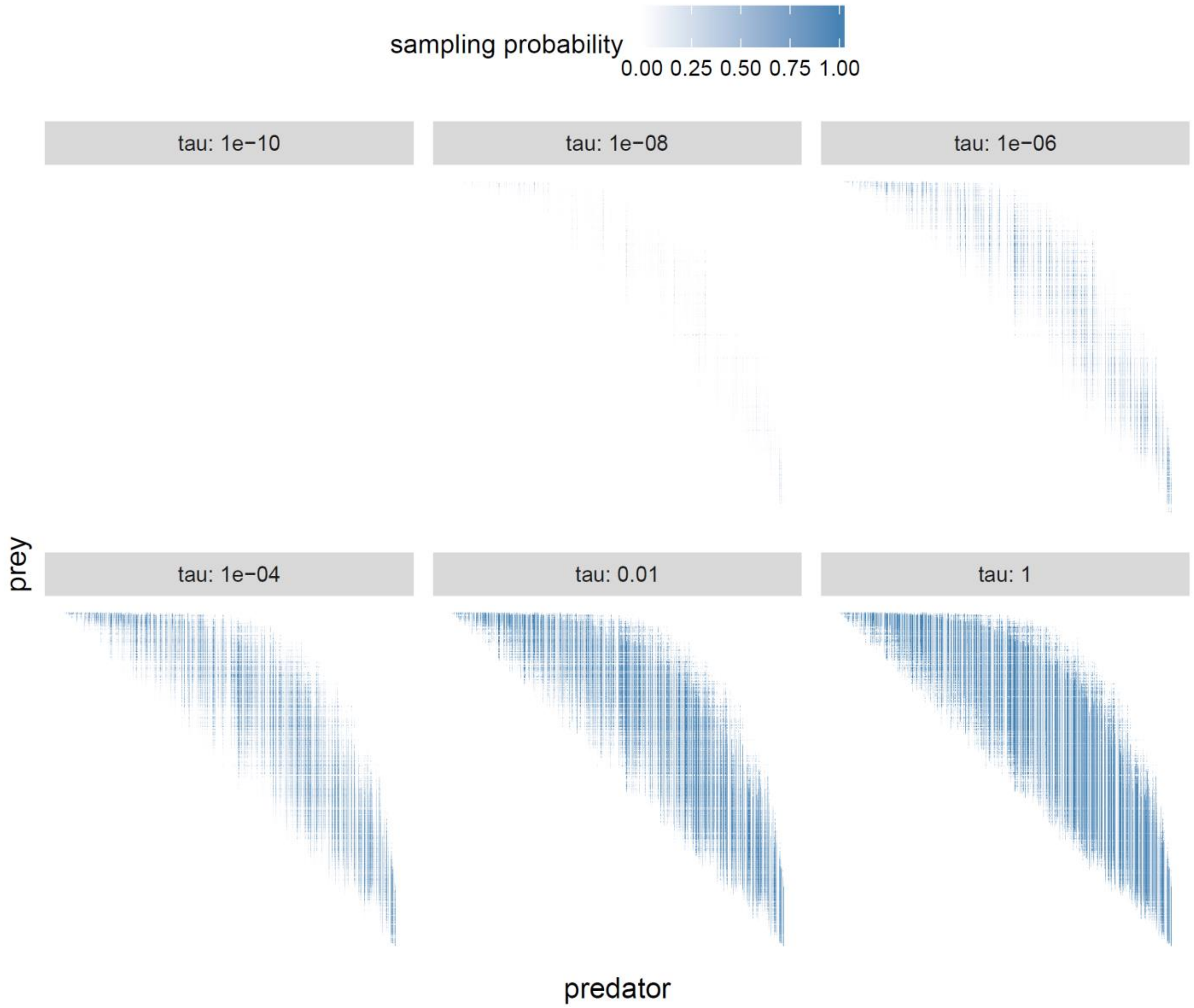


**Supplementary Figure 2: Effect of Sampling on the full Madingley Food web**. The figure shows the sampling probability of observing an interaction between preys and predators in the full Madingley food web output (without performing any cohort aggregation), represented as an adjacency matrix for different observation times (sampling effort) $\tau$'s. Rows (prey) and columns (predator) are sorted according to increase individual mass of each cohort.

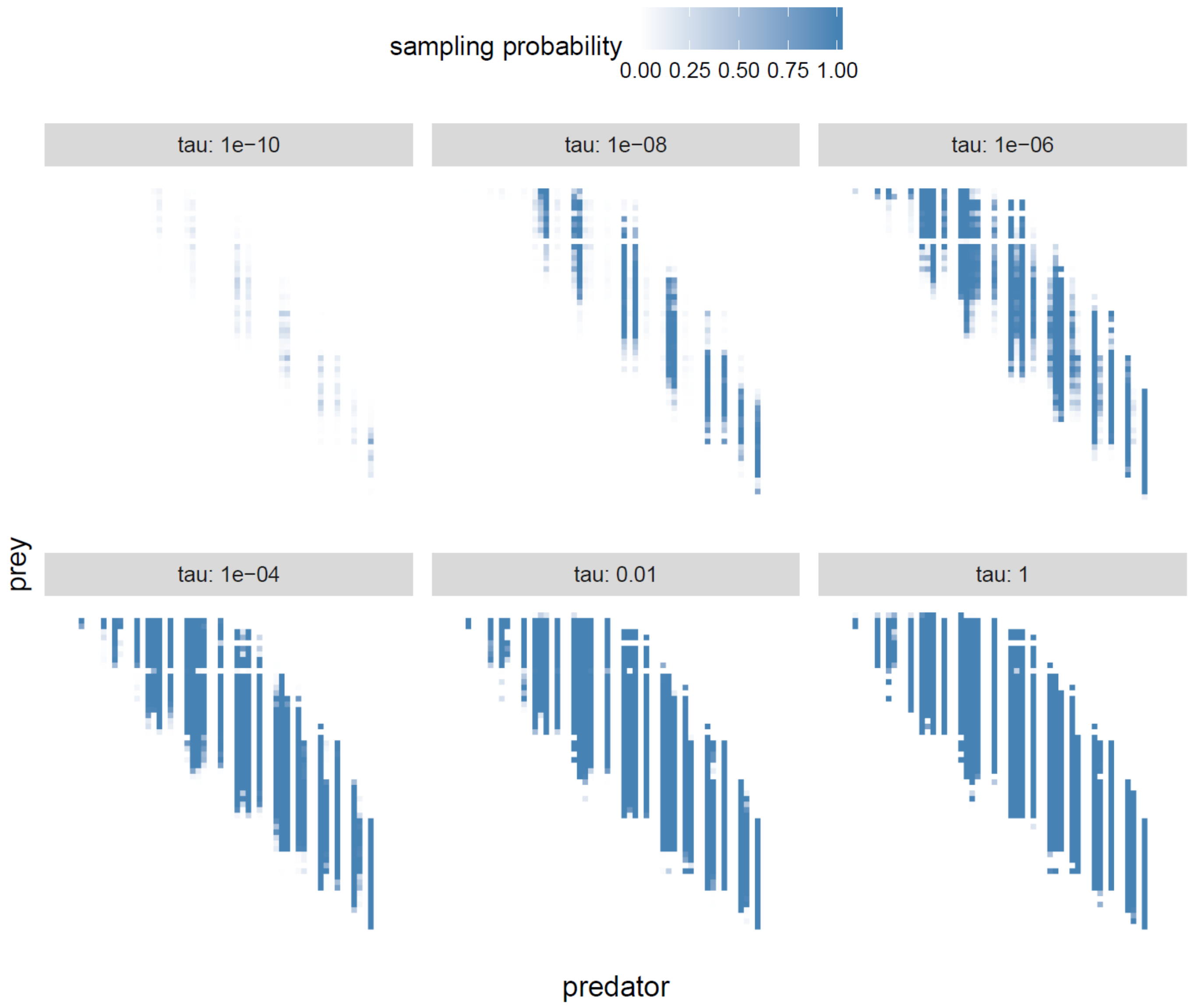


**Supplementary Figure 3: Effect of Sampling on an aggregated food web using Jaccard Trophic Similarity**.

The figure shows the sampling probability of observing an interaction between preys and predators on an aggregate Mandingley food web, represented as an adjacency matrix for different observation times (sampling effort) $\tau$'s. Cohorts were aggregated into $S$ = 60 different trophospecies using the **Jaccard** aggregation method. Rows (prey) and columns (predator) are sorted according to increase individual mass.

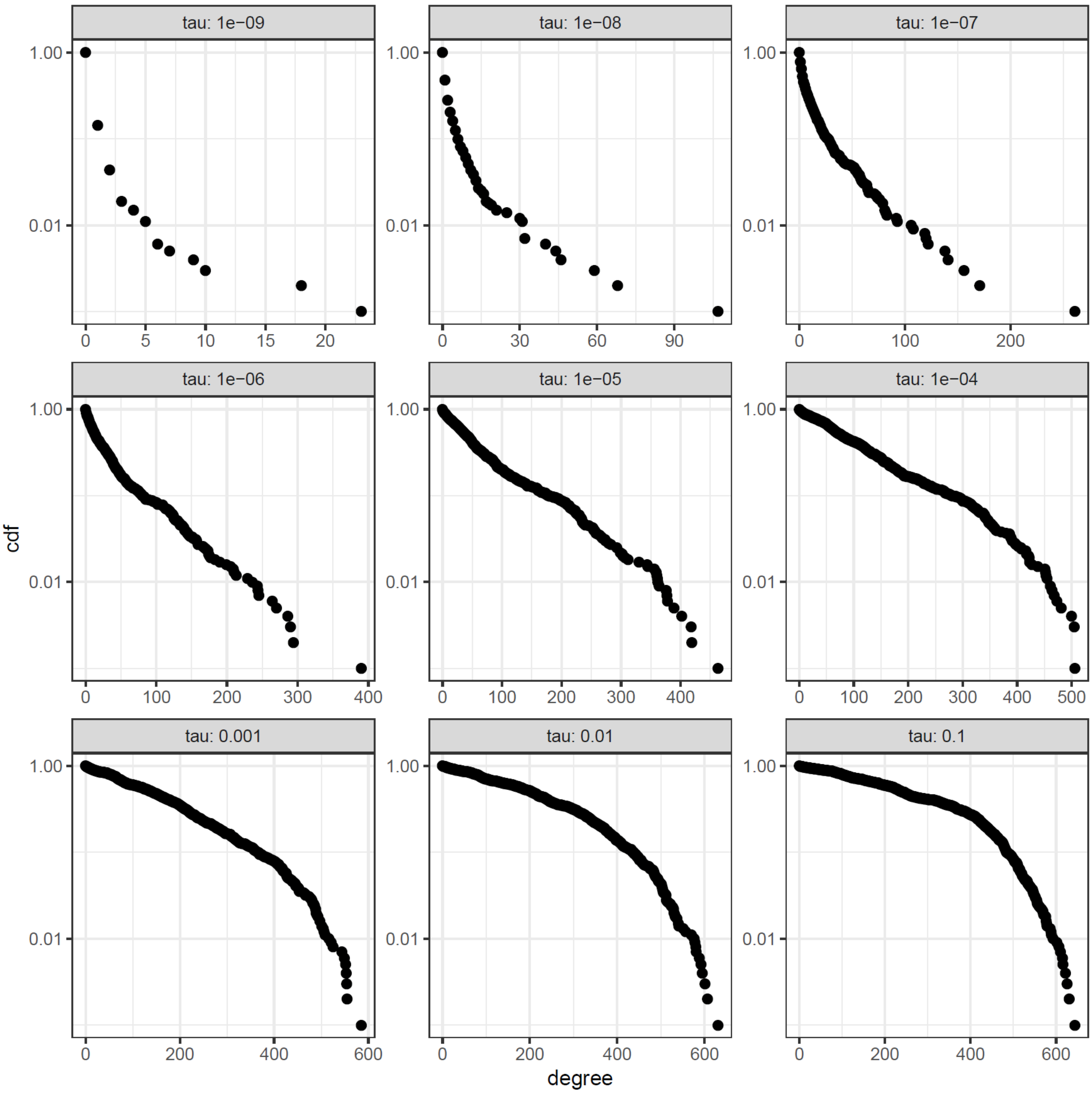


**Supplementary Figure 4: Effect of sampling on the degree distribution of the full Madingley food web**. The figure shows the expected degree distribution given observation time (sampling effort) $\tau$.

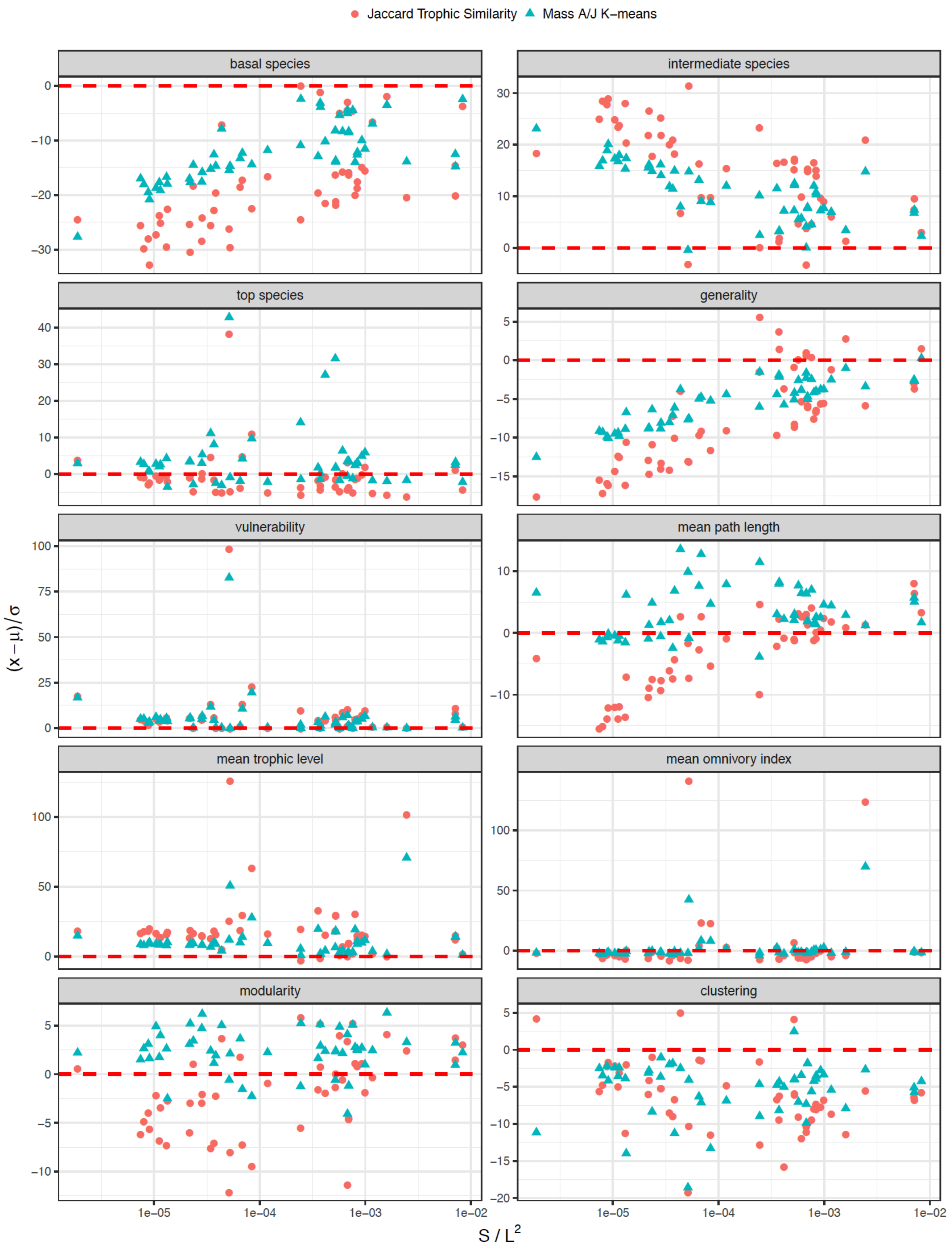


**Supplementary Figure 5: Comparison of Madingley sampled networks with empirical studies.** For each empirical study 1,000 aggregated sampled food webs were created with the same number of species and expected number of links (i.e. same expected connectance) by choosing the appropriate parameter sampling effort τ. Two data points (one for each aggregated method) exists for each empirical network. y-axis represent the ratio z-score of the absolute difference of empirical values, where μ and σ values come from the corresponding Madingley sampled networks.

**Supplementary Table 1:** Main description of the empirical food webs included in the present study

| *Reference* | *Ecosystem* | *Name* | *Species (S)* | *Links (L)* | *Connectance (C)* |
|---|---|---|---|---|---|
| *(Baird and Ulanowicz 1989)* | Estuary/salt marsh | Chesapeake bay | 36 | 121 | 0.09 |
| *(Jacob 2005, Brose et al. 2006a, Brose et al. 2006b)* | Marine | Weddell sea | 492 | 16136 | 0.07 |
| *(Cattin Blandenier 2004)* | Terrestrial | Grand Caricaie Cl C1 | 166 | 2080 | 0.08 |
| *(Cattin Blandenier 2004)* | Terrestrial | Grand Caricaie Cm C1 | 202 | 2930 | 0.07 |
| *(Cattin Blandenier 2004)* | Terrestrial | Grand Caricaie Cm M2 | 118 | 997 | 0.07 |
| *(Cattin Blandenier 2004)* | Terrestrial | Grand Caricaie Sn C2 | 152 | 1525 | 0.07 |
| *(Christian and Luczkovich 1999)* | Estuary/salt marsh | St. Mark's | 48 | 220 | 0.10 |
| *(Cohen 1989)* | Terrestrial | EcoWeb 59 | 30 | 65 | 0.07 |
| *(Cohen 1989)* | Terrestrial | EcoWeb 60 | 33 | 68 | 0.06 |
| *(Goldwasser and Roughgarden 1993)* | Terrestrial | St Martin | 44 | 218 | 0.11 |
| *(Harper-Smith et al. 2006)* | Freshwater | Sierra lakes | 37 | 298 | 0.22 |
| *(Harrison 2003)* | Freshwater | Alamitos creek | 162 | 3756 | 0.14 |
| *(Harrison 2003)* | Freshwater | Blackrock | 82 | 348 | 0.05 |
| *(Harrison 2003)* | Freshwater | Caldero creek | 126 | 2109 | 0.13 |
| *(Harrison 2003)* | Freshwater | Corde Matre creek | 106 | 1757 | 0.16 |
| *(Harrison 2003)* | Freshwater | Coyote | 190 | 4583 | 0.13 |
| *(Harrison 2003)* | Freshwater | Guadeloupe creek | 174 | 4662 | 0.15 |
| *(Harrison 2003)* | Freshwater | Guadeloupe river | 136 | 2487 | 0.13 |
| *(Harrison 2003)* | Freshwater | Los Gatos creek | 177 | 4480 | 0.14 |
| *(Harrison 2003)* | Freshwater | Los Trancos creek | 129 | 2440 | 0.15 |
| *(Harrison 2003)* | Freshwater | Penetetia creek | 170 | 4037 | 0.14 |
| *(Harrison 2003)* | Freshwater | San Francisquito creek | 140 | 3266 | 0.17 |
| *(Harrison 2003)* | Freshwater | Saratoga creek | 158 | 3754 | 0.15 |
| *(Harrison 2003)* | Freshwater | Steverson creek | 170 | 4776 | 0.17 |
| *(Havens 1992)* | Freshwater | Alford lake | 56 | 219 | 0.07 |
| *(Havens 1992)* | Freshwater | Balsam lake | 53 | 182 | 0.06 |
| *(Havens 1992)* | Freshwater | Beaver lake | 61 | 327 | 0.09 |
| *(Havens 1992)* | Freshwater | Bridge brook lake | 75 | 553 | 0.10 |
| *(Havens 1992)* | Freshwater | Chub pond | 65 | 417 | 0.10 |
| *(Havens 1992)* | Freshwater | Connery lake | 30 | 60 | 0.07 |
| *(Havens 1992)* | Freshwater | Hoel lake | 49 | 254 | 0.11 |
| *(Havens 1992)* | Freshwater | Little rock lake | 176 | 2009 | 0.06 |
| *(Havens 1992)* | Freshwater | Long lake | 65 | 416 | 0.10 |
| *(Havens 1992)* | Freshwater | Stink lake | 53 | 280 | 0.10 |
| *(Heymans et al. 2002)* | Estuary/salt marsh | Mangrove estuary | 94 | 1339 | 0.15 |
| *Jacob (unpub. data)* | Marine | Lough hyne | 350 | 5114 | 0.04 |
| *(Link 2002)* | Marine | NE US shelf | 81 | 1482 | 0.23 |
| *Navarette & Wieters (unpub data)* | Marine | Chile food web | 106 | 1436 | 0.13 |
| *(Patrício and Marques 2006)* | Marine | Mondego Zostera meadows | 47 | 278 | 0.13 |
| *(Polis 1991)* | Terrestrial | Coachella | 27 | 228 | 0.31 |
| *(Simberloff and Abele 1976)* | Terrestrial | Simberloff_E1 | 48 | 239 | 0.10 |
| *(Simberloff and Abele 1976)* | Terrestrial | Simberloff_E2 | 63 | 347 | 0.09 |
| *(Simberloff and Abele 1976)* | Terrestrial | Simberloff_E3 | 49 | 242 | 0.10 |
| *(Simberloff and Abele 1976)* | Terrestrial | Simberloff_E7 | 52 | 255 | 0.09 |
| *(Simberloff and Abele 1976)* | Terrestrial | Simberloff_E9 | 71 | 446 | 0.09 |
| *(Simberloff and Abele 1976)* | Terrestrial | Simberloff_ST2 | 63 | 347 | 0.09 |
| *(Townsend 1998)* | Freshwater | Broad 2 | 34 | 221 | 0.19 |
| *(Townsend 1998)* | Freshwater | Ross | 117 | 2024 | 0.15 |
| *(Waide and Reagan 1996)* | Terrestrial | El Verde | 156 | 1509 | 0.06 |
| *(Warren 1989)* | Freshwater | Skipwith pond | 35 | 379 | 0.31 |
| *(Woodward et al. 2005)* | Freshwater | Broadstone stream | 34 | 221 | 0.19 |
| *Woodward et al. (?)* | Freshwater | Bere stream | 137 | 1276 | 0.07 |

**Supplementary Table 2:** Network metrics of the empirical food webs included in the present study

| *Name* | *S* | *L* | *C* | *basal species* | *interm. species* | *top species* | *gen.* | *vul.* | *mean path length* | *mean trophic level* | *mean omnivory index* | *mod.* | *trans.* |
|---|---|---|---|---|---|---|---|---|---|---|---|---|---|
| *Alamitos creek* | 162 | 3756 | 0.14 | 6 | 151 | 5 | 24.08 | 23.92 | 2.08 | 2.62 | 0.23 | 0.21 | 0.46 |
| *Alford lake* | 56 | 219 | 0.07 | 24 | 31 | 1 | 6.84 | 3.98 | 1.44 | 1.70 | 0.02 | 0.26 | 0.18 |
| *Balsam lake* | 53 | 182 | 0.06 | 31 | 21 | 1 | 8.27 | 3.50 | 1.50 | 1.51 | 0.02 | 0.40 | 0.09 |
| *Beaver lake* | 61 | 327 | 0.09 | 27 | 34 | 0 | 9.62 | 5.36 | 1.58 | 1.75 | 0.05 | 0.33 | 0.19 |
| *Bere stream* | 137 | 1276 | 0.07 | 12 | 94 | 31 | 10.21 | 12.04 | 1.54 | 4.33 | 0.53 | 0.23 | 0.12 |
| *Blackrock* | 82 | 348 | 0.05 | 44 | 22 | 16 | 9.16 | 5.27 | 1.36 | 1.53 | 0.01 | 0.16 | 0.04 |
| *Bridge brook lake* | 75 | 553 | 0.10 | 39 | 36 | 0 | 15.36 | 7.37 | 1.45 | 1.64 | 0.04 | 0.36 | 0.19 |
| *Broad 2* | 34 | 221 | 0.19 | 5 | 28 | 1 | 7.62 | 6.70 | 1.45 | 2.12 | 0.10 | 0.08 | 0.47 |
| *Broadstone stream* | 34 | 221 | 0.19 | 5 | 28 | 1 | 7.62 | 6.70 | 1.45 | 2.12 | 0.10 | 0.08 | 0.47 |
| *Caldero creek* | 126 | 2109 | 0.13 | 6 | 115 | 5 | 17.58 | 17.43 | 2.21 | 2.60 | 0.24 | 0.23 | 0.41 |
| *Chesapeake bay* | 36 | 121 | 0.09 | 2 | 34 | 0 | 3.56 | 3.36 | 2.75 | 8.10 | 3.33 | 0.28 | 0.28 |
| *Chile food web* | 106 | 1436 | 0.13 | 7 | 50 | 49 | 14.51 | 25.19 | 1.33 | 2.93 | 0.14 | 0.16 | 0.08 |
| *Chub pond* | 65 | 417 | 0.10 | 33 | 32 | 0 | 13.03 | 6.42 | 1.55 | 1.70 | 0.05 | 0.38 | 0.19 |
| *Coachella* | 27 | 228 | 0.31 | 3 | 23 | 1 | 9.50 | 8.77 | 1.42 | 2.78 | 0.46 | 0.09 | 0.74 |
| *Connery lake* | 30 | 60 | 0.07 | 19 | 10 | 1 | 5.45 | 2.07 | 1.33 | 1.40 | 0.02 | 0.34 | 0.09 |
| *Corde Matre creek* | 106 | 1757 | 0.16 | 6 | 90 | 10 | 17.57 | 18.30 | 1.73 | 2.51 | 0.19 | 0.14 | 0.47 |
| *Coyote* | 190 | 4583 | 0.13 | 6 | 180 | 4 | 24.91 | 24.64 | 2.52 | 2.72 | 0.27 | 0.24 | 0.41 |
| *EcoWeb 59* | 30 | 65 | 0.07 | 7 | 15 | 8 | 2.83 | 2.95 | 1.46 | 2.27 | 0.06 | 0.28 | 0.07 |
| *EcoWeb 60* | 33 | 68 | 0.06 | 5 | 17 | 11 | 2.43 | 3.09 | 1.55 | 2.34 | 0.04 | 0.39 | 0.03 |
| *El Verde* | 156 | 1509 | 0.06 | 28 | 107 | 21 | 11.79 | 11.18 | 2.54 | 2.92 | 0.54 | 0.19 | 0.23 |
| *Grand Caricaie Cl C1* | 166 | 2080 | 0.08 | 21 | 144 | 1 | 14.34 | 12.61 | 2.66 | 2.51 | 0.16 | 0.27 | 0.20 |
| *Grand Caricaie Cm C1* | 202 | 2930 | 0.07 | 35 | 166 | 1 | 17.54 | 14.58 | 2.41 | 2.49 | 0.19 | 0.34 | 0.25 |
| *Grand Caricaie Cm M2* | 118 | 997 | 0.07 | 20 | 97 | 1 | 10.17 | 8.52 | 2.73 | 2.52 | 0.24 | 0.29 | 0.24 |
| *Grand Caricaie Sn C2* | 152 | 1525 | 0.07 | 23 | 126 | 3 | 11.82 | 10.23 | 2.49 | 2.52 | 0.25 | 0.30 | 0.25 |
| *Guadeloupe creek* | 174 | 4662 | 0.15 | 6 | 163 | 5 | 27.75 | 27.59 | 2.10 | 2.64 | 0.23 | 0.20 | 0.47 |
| *Guadeloupe river* | 136 | 2487 | 0.13 | 6 | 124 | 6 | 19.13 | 19.13 | 2.35 | 2.66 | 0.28 | 0.25 | 0.43 |
| *Hoel lake* | 49 | 254 | 0.11 | 22 | 27 | 0 | 9.41 | 5.18 | 1.62 | 1.83 | 0.07 | 0.34 | 0.24 |
| *Little rock lake* | 176 | 2009 | 0.06 | 62 | 113 | 1 | 17.62 | 11.48 | 1.66 | 2.00 | 0.08 | 0.37 | 0.32 |
| *Long lake* | 65 | 416 | 0.10 | 30 | 33 | 2 | 11.89 | 6.60 | 1.53 | 1.77 | 0.06 | 0.24 | 0.29 |
| *Los Gatos creek* | 177 | 4480 | 0.14 | 6 | 168 | 3 | 26.20 | 25.75 | 2.20 | 2.67 | 0.26 | 0.22 | 0.47 |
| *Los Trancos creek* | 129 | 2440 | 0.15 | 6 | 118 | 5 | 19.84 | 19.68 | 1.96 | 2.60 | 0.21 | 0.21 | 0.44 |
| *Lough hyne* | 350 | 5114 | 0.04 | 50 | 287 | 13 | 17.05 | 15.18 | 2.39 | 2.39 | 0.14 | 0.34 | 0.11 |
| *Mangrove estuary* | 94 | 1339 | 0.15 | 5 | 89 | 0 | 15.04 | 14.24 | 2.21 | 6.63 | 3.06 | 0.17 | 0.42 |
| *Mondego Zostera meadows* | 47 | 278 | 0.13 | 10 | 31 | 6 | 7.51 | 6.78 | 1.48 | 2.10 | 0.09 | 0.20 | 0.33 |
| *NE US shelf* | 81 | 1482 | 0.23 | 3 | 75 | 3 | 19.00 | 19.00 | 1.77 | 3.04 | 0.25 | 0.12 | 0.58 |
| *Penetetia creek* | 170 | 4037 | 0.14 | 6 | 158 | 6 | 24.62 | 24.62 | 2.26 | 2.65 | 0.24 | 0.22 | 0.46 |
| *Ross* | 117 | 2024 | 0.15 | 6 | 105 | 6 | 18.23 | 18.23 | 2.48 | 2.64 | 0.26 | 0.24 | 0.48 |
| *San Francisquito creek* | 140 | 3266 | 0.17 | 6 | 129 | 5 | 24.37 | 24.19 | 2.05 | 2.64 | 0.24 | 0.22 | 0.47 |
| *Saratoga creek* | 158 | 3754 | 0.15 | 6 | 148 | 4 | 24.70 | 24.38 | 2.05 | 2.66 | 0.25 | 0.22 | 0.46 |
| *Sierra lakes* | 37 | 298 | 0.22 | 4 | 29 | 4 | 9.03 | 9.03 | 1.37 | 2.34 | 0.16 | 0.15 | 0.46 |
| *Simberloff_E1* | 48 | 239 | 0.10 | 4 | 39 | 5 | 5.43 | 5.56 | 1.33 | 2.46 | 0.13 | 0.25 | 0.29 |
| *Simberloff_E2* | 63 | 347 | 0.09 | 4 | 54 | 5 | 5.88 | 5.98 | 1.56 | 3.32 | 0.15 | 0.28 | 0.28 |
| *Simberloff_E3* | 49 | 242 | 0.10 | 3 | 41 | 5 | 5.26 | 5.50 | 1.52 | 2.63 | 0.15 | 0.25 | 0.28 |
| *Simberloff_E7* | 52 | 255 | 0.09 | 2 | 45 | 5 | 5.10 | 5.43 | 1.59 | 3.50 | 0.16 | 0.25 | 0.27 |
| *Simberloff_E9* | 71 | 446 | 0.09 | 5 | 61 | 5 | 6.76 | 6.76 | 1.73 | 3.59 | 0.25 | 0.26 | 0.27 |
| *Simberloff_ST2* | 63 | 347 | 0.09 | 4 | 54 | 5 | 5.88 | 5.98 | 1.56 | 3.32 | 0.15 | 0.28 | 0.28 |
| *Skipwith pond* | 35 | 379 | 0.31 | 1 | 33 | 1 | 11.15 | 11.15 | 1.14 | 2.65 | 0.17 | 0.04 | 0.62 |
| *St Martin* | 44 | 218 | 0.11 | 8 | 30 | 6 | 6.06 | 5.74 | 1.64 | 2.58 | 0.19 | 0.25 | 0.33 |
| *St. Mark's* | 48 | 220 | 0.10 | 7 | 32 | 9 | 5.37 | 5.64 | 1.79 | 2.61 | 0.22 | 0.20 | 0.29 |
| *Steverson creek* | 170 | 4776 | 0.17 | 6 | 159 | 5 | 29.12 | 28.95 | 2.11 | 2.74 | 0.30 | 0.20 | 0.50 |
| *Stink lake* | 53 | 280 | 0.10 | 24 | 28 | 1 | 9.66 | 5.38 | 1.43 | 1.71 | 0.03 | 0.33 | 0.19 |
| *Weddell sea* | 492 | 16136 | 0.07 | 70 | 389 | 33 | 38.24 | 35.15 | 2.20 | 2.59 | 0.21 | 0.29 | 0.26 |

Baird, D., and R. E. Ulanowicz. 1989. The Seasonal Dynamics of The Chesapeake Bay Ecosystem. Ecological Monographs **59**:329-364.

Brose, U., T. Jonsson, E. Berlow, P. Warren, C. Banasek-Richter, L.-F. Bersier, J. L Blanchard, T. Brey, S. Carpenter, M.-F. Cattin Blandenier, L. Cushing, H. Dawah, T. Dell, F. Edwards, S. Harper-Smith, U. Jacob, M. Ledger, N. Martinez, J. Memmott, and J. Cohen. 2006a. Consumer-resource body-size relationships in natural food webs.

Brose, U., R. Williams, and N. Martinez. 2006b. Allometric scaling enhances stability in complex food webs.

Cattin Blandenier, M.-F. 2004. Food web ecology: models and application to conservation. Thesis, University of Neuchatel, Switzerland. PhD. Université de Neuchâtel.

Christian, R. R., and J. J. Luczkovich. 1999. Organizing and understanding a winter's seagrass foodweb network through effective trophic levels. Ecological Modelling **117**:99-124.

Cohen, J. 1989. Ecologists' co-operative web bank, version 1.0. New York, NY: Rockefeller University.

Goldwasser, L., and J. Roughgarden. 1993. Construction and Analysis of a Large Caribbean Food Web. Ecology **74**:1216-1233.

Harper-Smith, S., E. L. Berlow, R. A. Knapp, R. J. Williams, and N. D. Martinez. 2006. Communicating ecology through food webs: visualizing and quantifying the effects of stocking alpine lakes with trout. Dynamic Food Webs. Elsevier Inc.

Harrison, K. A. 2003. Effects of Land Use and Dams on Stream Food Web Ecology in Santa Clara Valley, California. San Francisco State University.

Havens, K. 1992. Scale and Structure in Natural Food Webs. Science **257**:1107-1109.

Heymans, J. J., R. E. Ulanowicz, and C. Bondavalli. 2002. Network analysis of the South Florida Everglades graminoid marshes and comparison with nearby cypress ecosystems. Ecological Modelling **149**:5-23.

Jacob, U. 2005. Trophic Dynamics of Antarctic Shelf Ecosystems - Food Webs and Energy Flow Budgets. PhD. Universität Bremen.

Link, J. 2002. Does food web theory work for marine ecosystems? Marine Ecology Progress Series **230**:1-9.

Patrício, J., and J. C. Marques. 2006. Mass balanced models of the food web in three areas along a gradient of eutrophication symptoms in the south arm of the Mondego estuary (Portugal). Ecological Modelling **197**:21-34.

Polis, G. A. 1991. Complex Trophic Interactions in Deserts: An Empirical Critique of Food-Web Theory. The American Naturalist **138**:123-155.

Simberloff, D. S., and L. G. Abele. 1976. Island Biogeography Theory and Conservation Practice. Science **191**:285-286.

Townsend, C. 1998. Disturbance, resource supply, and food-web architecture in streams. Ecology letters **1**:200-209.

Waide, R., and W. Reagan. 1996. The food web of a tropical rainforest. University of.

Warren, P. H. 1989. Spatial and temporal variation in the structure of a freshwater food web. Oikos:299-311.

Woodward, G., D. C. Speirs, A. G. Hildrew, and C. Hal. 2005. Quantification and resolution of a complex, size-structured food web. Advances in ecological research **36**:85-135.
